# A cassava drought inducible CC-type glutaredoxin, *MeGRX232*, negatively regulates drought tolerance in *Arabidopsis* by inhibition of ABA-dependent stomatal closure

**DOI:** 10.1101/125385

**Authors:** Meng-Bin Ruan, Yi-Ling Yang, Xin Guo, Xue Wang, Bin Wang, Xiao-Ling Yu, Peng Zhang, Ming Peng

## Abstract

CC-type glutaredoxins (GRXs) are a land plant-specific GRX subgroup that evolved from CGFS GRXs, and participate in organ development and stress responses through the regulation of transcription factors. Here, genome-wide analysis identified 18 CC-type GRXs in the cassava genome, of which six (*MeGRX058, 232, 360, 496, 785*, and *892*) were induced by drought and ABA stress in cassava leaves. Furthermore, we found that overexpression of *MeGRX232* results in drought hypersensitivity in soil-grown plants, with a higher water loss rate, but with increased tolerance of mannitol and ABA in *Arabidopsis* on the sealed agar plates. The ABA induced stomatal closure is impaired in *MeGRX232-OE Arabidopsis*. Further analysis reveals that the overexpression of *MeGRX232* leads to more ROS accumulation in guard cells. MeGRX232 can interact with TGA5 from *Arabidopsis* and MeTGA074 from cassava *in vitro* and *in vivo*. The results of microarray assays show that *MeGRX232-OE* affected the expression of a set of drought and oxidative stress related genes. Taken together, we demonstrated that CC-type GRXs involved in ABA signal transduction and play roles in response to drought through regulating stomatal closure.

**Novelty statement:** We found that drought and ABA stress induced the transcription of CC-type glutaredoxins (GRXs) in cassava leaves. Ectopic expression of one of them, *MeGRX232* in *Arabidopsis* affected the sensitivity to abscisic acid (ABA) and mannitol, and caused drought hypersensitivity by impairment of ABA-dependent stomatal closure.

## Introduction

As a tropical crop, cassava (*Manihot esculenta*) evolved different response to drought stress, such as quick stomata closure, reduction of photosynthetic proteins levels and photosynthetic capacity, induction of senescence in older leaves, and size reduction of leave epidermal cells (Alves and Setter 2004; Zhao *et al*. 2014). Some cassava cultivars display faster senescence in older leaves than the others (Zhao *et al*. 2014). Senescence in cassava is partly controlled by a reactive oxygen species (ROS) and ethylene signaling (Liao *et al*. 2016b). Increasing the ROS scavenging ability in cassava delay leaf senescence under drought stress (Liao *et al*. 2016b; Xu *et al*. 2013a, 2013b). It is therefore necessary to analyze genes involved in these pathways for a deeper functional characterization.

Glutaredoxin (GRX) is one of the most important protein modification system in plants (Rouhier *et al*. 2006). The glutathione/GRX (GSH/GRX) system is essential for redox homeostasis and ROS signal in plant cells (Meyer *et al*. 2012). GRX target proteins are involved in all aspects of plant growth, including basal metabolism, iron/sulfur cluster formation, development, adaptation to the environment, and stress responses (Meyer *et al*. 2012). GRX are in particular studied for their involvement in oxidative stress responses (Carroll *et al*. 2006; Kanda *et al*. 2006; Meyer *et al*. 2012). GRXs are classified in five subgroups, among which CC-type GRXs are a plant-specific subgroup, also known as the ROXY family in *Arabidopsis* (Xing *et al*. 2005; Ziemann *et al*. 2009). CC-type GRXs likely evolved from the CPYC subgroup and expanded during land plant evolution (Ziemann *et al*. 2009). There are only two CC-type GRXs in the basal land plant *Physcomitrella*, but between 15 and 24 members in land plants such as rice, *Arabidopsis, Vitis* and *Populus* (Ziemann *et al*. 2009). However, the number of CC-type GRXs in cassava remains unclear.

CC-type GRXs are characterized by the presence of a redox site CC*(C/S/G) as well as their disulfide reductase activity that uses glutathione as cofactor (Couturier *et al*. 2010). The first CC-type GRX has been identified as a regulator of petal development (Xing *et al*. 2005). However, CC-type GRXs are also involved in jasmonic acid (JA)/ethylene mediated biotic stress responses through the interaction with TGA factors in *Arabidopsis* (La Camera *et al*. 2011; Wang *et al*. 2009; Zander *et al*. 2012). Moreover, some CC-type GRXs are critical in limiting basal and photo-oxidative stress-induced ROS production (Laporte *et al*. 2012). Thus, CC-type GRXs may play a key role in the crosstalk between ROS and ethylene. CC-type GRX members are also involved in organ development and biotic stress responses in other plants (El-Kereamy *et al*. 2015; Garg *et al*. 2010; Gutsche *et al*. 2015; Hong *et al*. 2012; R. Sharma *et al*. 2013; Wang *et al*. 2009).

During evolution, CC-type GRXs might have gained new functions in high land plants (Rouhier *et al*. 2006; Wang *et al*. 2009; Ziemann *et al*. 2009). Although several CC-type GRXs have been characterized in *Arabidopsis* and rice (Gutsche *et al*. 2015; Wang *et al*. 2009), no previous work profiled them in cassava. As a tropical crop, cassava evolved to be tolerant to intermittent drought, but hypersensitive to cold (An *et al*. 2012; Xia *et al*. 2014; Zeng *et al*. 2014; Zhao *et al*. 2014). Genome-wide analysis base on high-quality sequencing data and EST predict the presence of many genes related to abiotic stress responses in cassava (An *et al*. 2012; W. Hu *et al*. 2015b; W.Hu *et al*. 2015a; Wei Hu *et al*. 2016; Lokko *et al*. 2007; Sakurai *et al*. 2007; Xia *et al*. 2014; Zeng *et al*. 2014; Zhao *et al*. 2014). Therefore, it is now possible to analyze the expression pattern of a whole gene family to identify drought stress related members, and to characterize their functions during abiotic stresses response (W.Hu *et al*. 2015b; W.Hu *et al*. 2015a; Wei Hu *et al*. 2016; Xia *et al*. 2014; Zeng *et al*. 2014; Zhao *et al*. 2014).

In this study, we performed computational and phylogenetic analyses to identify plant specific CC-type GRXs in the cassava genome. In total, we identified 18 CC-type GRXs in cassava. Due to the lack of characterization of the function of CC-type GRXs during drought responses in cassava, we chose this subgroup for systematic and functional analysis. Based on our previously reported transcriptomic data of cassava cultivars (Wei Hu *et al*. 2016), we identified six CC-type GRXs (*MeGRX058, 232, 360, 496, 785, 892*) that responded to drought using qPCR analysis in cassava leaves. To investigate potential CC-type GRX-mediated drought stress responsive pathways, we examined the expression levels of these six genes in leaves under ABA treatment. Our results showed that CC-type GRXs may functions as component in drought stress in an ABA-dependent pathway in both Arg7 and SC124 plants. Overexpression of *MeGRX232* induces insensitivity to ABA and mannitol, and confers drought susceptibility in *Arabidopsis* in soil-grown condition by inhibiting ABA-dependent stomatal closing. The overexpression of *MeGRX232* caused more ROS accumulation in guard cells. In addition, gene expression analysis reveals that MeGRX232 regulated a set of oxidative stress related genes in *Arabidopsis*.

## Materials and Methods

### Plant materials

Stems of cassava Arg7 and SC124 were cultured in pots (36 cm in diameter x 30 cm in height) containing well-mixed soil (soil: vermiculite: pellets, 1:1:1) for 80 days in greenhouse at the Institute of Tropical Bioscience and Biotechnology (HaiKou, China). Wild type (Col-0) *Arabidopsis* plants for transformation were grown in 12 hrs light/12 hrs dark at 20-23 □ until the primary inflorescence was 5-15 cm tall and a secondary inflorescence appeared at the rosette.

### Drought and ABA treatments

For drought treatment, watering was interrupted for 14 days. Leaves were collected from three Arg7 and SC124 plants eight or 14 days from the start of the drought treatment and 24 hours after re-water at the end of the treatment. Plants watered as normal were used as control. Three leaves (the second, third and fourth leaf from the top of plant) from each plant were collected. For ABA treatment, mature leaves with petiole were excised from Arg7 and SC124 plants, treated by dipping the petioles in water with 20 μM ABA respectively for 30 min before collection.

### Bioinformatics analysis

The protein sequences of cassava GRXs were predicted using a TBLASTN search against the cassava genome database in Phytozome (https://phvtozome.jgi.doe.gov, *Manihot esculenta v4.1*) with the protein sequence from *Arabidopsis* GRXs as a query. All *Arabidopsis* GRX protein sequences were downloaded from GenBank (Supplementary Table S1). Multiple sequence alignments were conducted using ClustalW (Thompson *et al*. 1994). An unrooted phylogenetic tree showing cassava GRXs and *Arabidopsis* GRX family was generated via the neighbor joining method using MEGA5.0 (Tamura *et al*. 2011). Editing of aligned sequences of cassava CC-type GRXs was performed using AlignX (Vector NTI suite 10.3, Invitrogen).

### Transcriptome data analysis

For drought responsive CC-type GRXs identification, we used our previously reported RNA-seq data (Wei Hu *et al*. 2016). We used data that included two tissues (leaf and root) under drought treatment (12 day after water withholding) and a control. Gene expression levels were normalized using FPKM. We selected CC-type GRX genes, and generated a heat map and hierarchical clustering using Cluster 3.0. RNA-seq datasets are available at NCBI and the accession numbers are listed in Supplementary Table S2.

### Quantitative real-time PCR (qPCR) analysis

Total RNA was isolated from leaves of cassava plant used a modified CTAB method. cDNA synthesis was performed with FastQuant RT Kit (TIANGEN). Expression analysis of CC-type GRXs in cassava leaves after drought and exogenous ABA treatment were performed by qPCR with gene-specific primers (Supplementary Table S3). For qPCR analysis in transgenic plants, total RNA was isolated from wild type, three independent *MeGRX785-OE* and *MeGRX058-OE* transgenic lines, respectively. All qPCR reactions were carried out in triplicates, with SYBR^®^ Premix Ex Taq^TM^ II Kit (Takara) on StepOne™ Real-Time PCR system (Applied Biosystems), and the comparative ΔΔCT method employed to evaluate amplified product quantities in the samples.

### DNA constructs, protein subcellular localization, and Arabidopsis transformation

Full-length coding sequence without stop-codon of *MeGRX232* was isolated from cDNA of drought stressed leaves by RT-PCR. Fragments were identified by sequencing and fused to *GFP* behind the *CaMV 35S* promoter in the modified plant expression vector *pG1300* (*eGFP:pCAMBIA1300*) to make *P35s:MeGRX232:GFP*,. The *P35s:MeGRX232:GFP* and vector construct were transferred into *Agrobacterium LBA4404* respectively. Leaves from four-week-old *Nicothiana benthamiana* plants were transformed by infiltration of *Agrobacterium* cells (OD_600_=1.2) harboring appropriate DNA constructs using 5-mL syringe without needle. The *pG1300* vector (*GFP*) and *P35s:MeHistone3:GFP* (*H3:GFP*) were used as the positive controls. After three days, infiltrated *N.benthamiana* leaves were imaged for reconstitution of GFP fluorescence by confocal laser scanning microscope (Olympus FluoView FV1100). *Arabidopsis* was transformed using the DIP method (Clough and Bent 1998) with *A. tumefaciens* strain *LBA4404* carrying the DNA constructs *P35s:MeGRX232:GFP*. More than three homozygote lines of each construct were selected for further phenotype analysis.

### ABA and mannitol stress tolerance assays of transgenic Arabidopsis

To study the response of *MeGRX232-OE* transgenic plants to ABA, 5-d-old seedlings were transferred to MS medium contained with 0 μM (mock) and 5 μM ABA grown for 10 days. For mannitol treatment, 5-d-old seedlings were transferred to MS medium contained with 0 mM (mock) and 250 mM D-mannitol grown for 15 days. Rosette diameter, primary root length and later root number were measured. The transgenic plant that contained *pG1300* vector was used as control.

### Drought stress tolerance assays of transgenic Arabidopsis

Post-germinated seedlings of *MeGRX232-OE* and vector transgenic plants were grown in soil in one pot for 15 days under normal conditions. For drought stress, the plants were treated by water withholding for 21 days, then re-watering. Survival rates have been calculated at 5 days after re-watering. Lipid peroxidation in transgenic *Arabidopsis* leaves was measured in terms of MDA in the samples as described in reference (Xu *et al*. 2013b) during drought stress. For water loss rate measurement, excised leaves from 28-d-old unstressed transgenic plants were kept on filter paper at room temperature. Their weight was measured after every 1 hour, up to 7 hour followed by calculation of water loss percentage.

### Stomatal aperture assays

ABA-induced stomatal closing assays were performed using fully expanded healthy leaves from 28-d-old transgenic plants as previous report (G. Sharma *et al*. 2015). Excised leaves were incubated in stomatal opening solution (10mM KCl, 100μM CaCl2, and 10mM MES, pH 6.1) for 2 hours followed by incubation in stomatal opening solution supplemented with varying ABA concentrations (0uM, 0.1uM, 1uM, 10uM) for 2 hrs. With the help of forceps, epidermal peels from the abaxial surface of treated leaves were peeled off and mounted on glass slides covered with coverslips followed by observation under Zeiss Scope A1 Imaging System. The ratio of width and length of stomata was measured using ZEN software. Approximately 30 guard cells were taken into account in measuring aperture in each sample.

### Determination of ROS accumulation

H_2_O_2_ was visualized by staining with DAB according to the reference (Thordal-Christensen *et al*. 1997). The untreated leaves of *MeGRX232-OE* and vector *Arabidopsis* were infiltrated with 2 mL of DAB solution (1mg mL^-1^ DAB, pH 3.8) in Eppendorf tube for 12 hours. Then the leaves were immersed in 95% (w/v) boiling ethanol for 10 min to decolorize the chloroplast. For the ROS accumulation assay in guard cells, prepared epidermal peels with or without H_2_O_2_, ABA treatment were load with 50 pM 2,7-dichlorofluorescin diacetate (DCFH-DA; Beyotime Biotechnology) for 30 min. The fluorescence was recorded by confocal microscopy with excitation at 546 nm. The fluorescence intensity was calculated from at least 30 guard cells by FV10-ASW (Olympus).

### Transactivation analysis and yeast two hybrid assay

Before analyzing the interaction between MeGRX232 and TGA factors, an autonomous transactivation analysis has been performed in yeast strain Y187. The MeGRX232 was in frame fused to GAL4 BD (binding domain) in *pGBKT7*, and then transformed into yeast Y187. Because MeGRX232 shows “autonomous transactivation” in yeast, a MeGRX232 GSH binding site mutant *MeGRX232mP_65_G_75_* was produced by replacing P_65_AVFIGGILVG_7_5 to A_65_AVFIGGILVA_75_. Next, for identification the interaction between MeGRX232 and TGA factors, a yeast two-hybrid assay has been performed in yeast strain Y187 based on the Matchmaker ^TM^ GAL4 two-hybrid system 3 manual (Clontech). The *MeGRX232* GSH binding site mutant DNA construct *MeGRX232mP_65_G_75_:pGBKT7* was used as bait. The cDNA sequences of TGA factors from *Arabidopsis* and cassava were introduced into the pGADT7, respectively in frame fused to GAL4 activate domain (AD). The *MeGRX232mP_65_G_75_:pGBKT7* and *TGA:pGADT7* constructs were pairwise co-transformed into yeast strain Y187. The presence of transgenes was confirmed by growth on SD/ -Trp/-Leu plates. Interactions between two proteins were checked by examining *β*-galactosidase activity as the manual instructed.

### Bimolecular fluorescence complementation analysis

To confirm the interactions between MeGRX232 and TGA2/MeTGA074 factors, a bimolecular fluorescence complementation assay has been performed by *N.benthamiana* transient system as previously report (Pazmino *et al*. 2011). The full-length coding sequence without stop-codon of *MeGRX232* was in frame fused to N- or C-terminus to yellow fluorescent protein (YFP) fragments (YN/YC) respectively to produce *Pro35S:MeGRX232:YN:pBiFC* and *Pro35S:MeGRX232:YC:pBiFC*. The full-length coding sequence without stop-codon of *TGA2* and *MeTGA074* were in frame fused to YC or YN respectively to produce *Pro35S:TGA2:YC:pBiFC, Pro35S:TGA2:YN:pBiFC, Pro35S:MeTGA074:YC:pBiFC*, and *Pro35S:MeTGA074:YN:pBiFC*. The resulting constructs were then introduced into *A. tumefaciens LBA4404* strains. Constructs were pair-wise transiently expressed in epidermal cells of tobacco leaves. 3-5 days after transient co-expression of protein pairs, reconstitution of YFP fluorescence was examined by confocal microscopy using GFP filter. Then the assays were performed as the method of proteins subcellular localization described. As positive controls, green fluorescent protein (eGFP) was tagged to the C-terminus of TGA2 and MeTGA074 respectively, transiently expressed in tobacco leaves.

### Microarray analysis of transgenic *Arabidopsis*

Microarray experiments were conducted using Affymetrix Arabidopsis ATH1 Genome Array. Experiments were performed as three biological repeats using cDNAs prepared independently from three individual homozygote lines of *MeGRX323-OE Arabidopsis* that have been phenotypic analyzed in plant growth. The transgenic *Arabidopsis* plants that carried the *pG1300* vector were used as control. The experiments and data analysis were performed under the instruction of Affymetrix. Total microarray data have been deposited in the NCBI GEO database with the accession number: GSE81136. Gene ontology (GO) analyses for significant enrichments of various categories (Supplementary Table S4) were performed using MAS 3.0 (http://bioinfo.capitalbio.com/mas3/). The Venn diagram is created by online tool (http://bioinformatics.psb.ugent.be/webtools/Venn/).

## Results

### Phylogenetic, gene structure and conserved motif analysis of cassava CC-type GRXs

We predicted a total of 38 putative GRX proteins in the cassava genome using the *Arabidopsis* ROXYs in a BLAST search against the genome of the cassava cultivar AM560 (https://phytozome.jgi.doe.gov, *Manihot esculenta v4.1*). To understand the relationship between GRX proteins in cassava and *Arabidopsis*, we built a neighbor-joining phylogenetic tree using MEGA5.0 on the basis of the protein sequences. The results showed that many cassava GRXs were highly similar to their *Arabidopsis* counterparts (Fig. 1). We found that the CC-type subgroup had the most members among the GRX subgroups in cassava. All putative cassava GRXs are classified in five subgroups as in *Arabidopsis* and rice (Meyer *et al*. 2012; Rouhier *et al*. 2006). CC-type GRXs are plant-specific GRXs, derived from the CPYC subgroup and expanded from basal to high land plant (Ziemann *et al*. 2009). Our result also demonstrated that all cassava CC-type GRXs evolved from three cassava CPYC GRXs (Fig. 1). Our analysis predicted 18 full-length CC-type GRX members of cassava (Table. 1), less than 21 of *Arabidopsis* (Xing and Zachgo 2008). Cassava CC-type GRX genes we represent on nine chromosomes (Table. 1) and generally possess no intron (Fig.2A). Almost all of cassava CC-type GRXs shared a redox site in N-terminus, an L**LL protein binding motif, and an ALWL motif at the C-terminus. However, two members, MeGRX954 and MeGRX956, did not present ALWL motif at C-terminus (Fig.2B). CC-type GRXs have a distinctive conserved CC(M/L)(C/S) redox site motif in *Arabidopsis*, whereas this motif is extends to C(C/G/F/Y/P)(M/L)(C/S/I/A) in rice (Rouhier *et al*. 2006; Wang *et al*. 2009; Xing and Zachgo 2008; Ziemann *et al*. 2009). Most of cassava CC-type GRXs shared a distinctive CCM(C/S) redox site (Fig.2B). However, this motif (CDMC) was extended in two CC-type GRXs (MeGRX785 and MeGRX892) in cassava.

**Figure 1.**
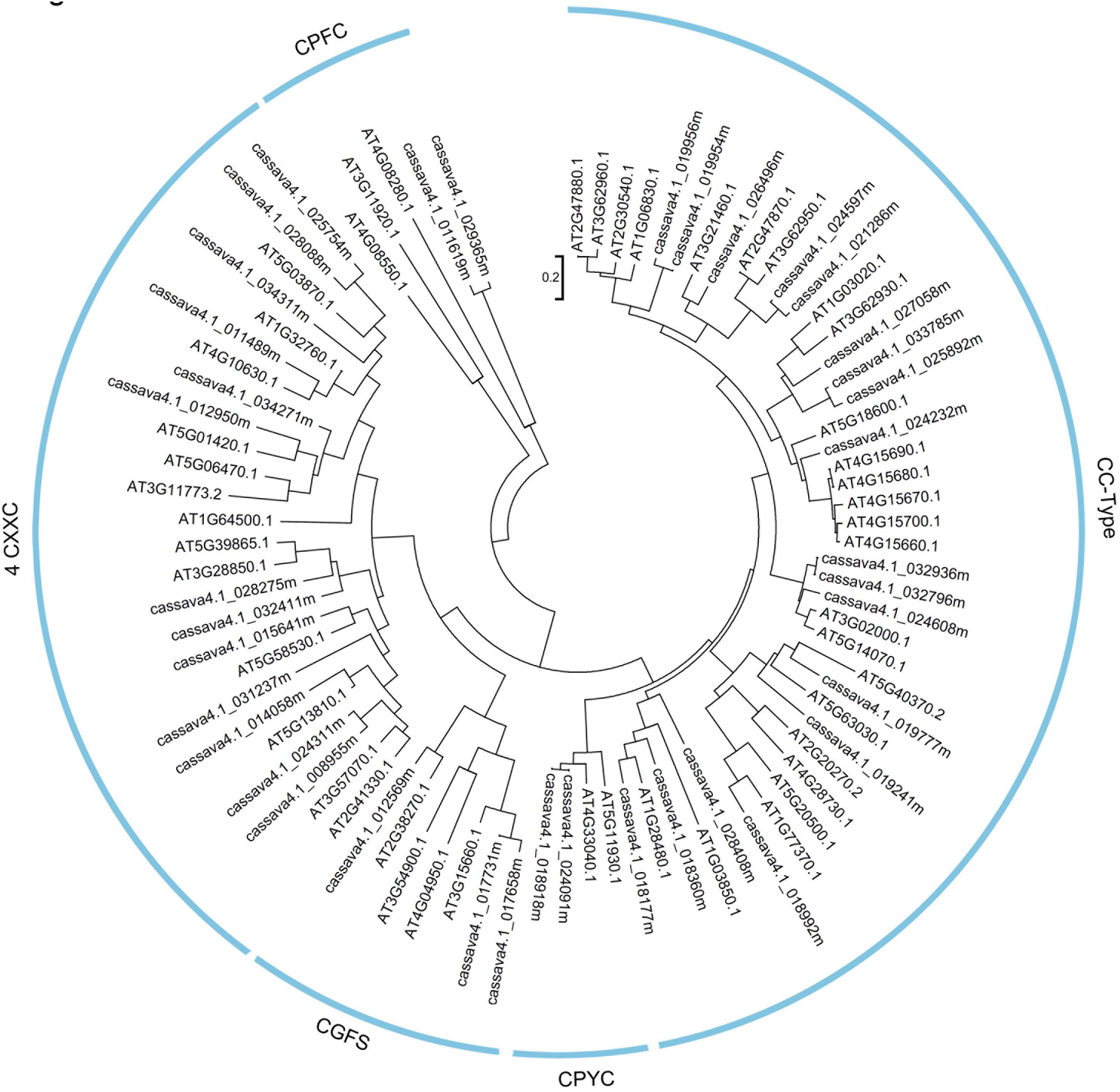
Phylogenetic tree of glutaredoxins from cassava and *Arabidopsis*. Multiple sequence alignments were conducted using the ClustalW program. An unrooted phylogenetic tree showing cassava GRXs and *Arabidopsis* GRXs was generated using the neighbor joining method using MEGA5.0. Members of GRXs were classified by their redox activate site. CC-type: CC-type subgroup; CPYC: CPYC subgroup; 4 CXXC: 4×CXXC subgroup; CGFS: CGFS subgroup; CPFC: CPFC subgroup.

**Table 1.** CC-type glutaredoxins in cassava

| Table 1. CC-type glutaredoxins in cassava |  |  |  |  |
| --- | --- | --- | --- | --- |
| JGI identifier (V4.1) | JGI identifier (V6.1) | Gene Symbol | Annotation | Chromosome location |
| cassava4.1_018177m | *Manes.13G141400.1* | *MeGRX177* | MeGRXS13 | Chromosome13:26980309..26981228 |
| cassava4.1_018360m | *Manes.17G050200.1* | *MeGRX360* | MeGRXC9 | Chromosome17:18791502..18792218 |
| cassava4.1_018918m | *Manes.03G049400.1* | *MeGRX918* | MeGRXC10-related | Chromosome03:4265637..4266041 |
| cassava4.1_019954m | *Manes.05G067000.1* | *MeGRX954* | MeGRXC13-related | Chromosome05:5123236..5123953 |
| cassava4.1_019956m | *Manes.01G214700.1* | *MeGRX956* | MeGRXC13-related | Chromosome01:30392540..30393324 |
| cassava4.1_021286m | *Manes.05G066900.1* | *MeGRX286* | MeGRXC11-related | Chromosome05:5120724..5122777 |
| cassava4.1_024091m | *Manes.16G081400.1* | *MeGRX091* | MeGRXC10-related | Chromosome16:23848065..23848499 |
| cassava4.1_024232m | *Manes.15G015500.1* | *MeGRX232* | MeGRXS2-related | Chromosome15:1265181..1265486 |
| cassava4.1_024597m | *Manes.01G214800.1* | *MeGRX597* | MeGRXC11-related | Chromosome01:30394902..30395210 |
| cassava4.1_024608m | *Manes.11G083800.1* | *MeGRX608* | MeGRXC7-related | Chromosome11:11464621..11464992 |
| cassava4.1_025892m | *Manes.05G066700.1* | *MeGRX892* | MeGRXS1-related | Chromosome05:5107836..5108531 |
| cassava4.1_026496m | *Manes.05G015600.1* | *MeGRX496* | MeGRXS10 | Chromosome15:1268904..1269746 |
| cassava4.1_027873m | *Manes.03G192100.1* | *MeGRX873* | MeGRXS2-related | Chromosome03: 27589330..27589919 |
| cassava4.1_027058m | *Manes.01G215100.1* | *MeGRX058* | MeGRXS1-related | Chromosome01:30426232..30427641 |
| cassava4.1_028408m | *Manes.15G124000.1* | *MeGRX408* | N/A | Chromosome15:9399566..9399949 |
| cassava4.1_032796m | *Manes.13G007200.1* | *MeGRX796* | MeGRXC7-related | Chromosome13:771539..771955 |
| cassava4.1_032936m | *Manes.12G007000.1* | *MeGRX936* | MeGRXC7-related | Chromosome12:652588..653004 |
| cassava4.1_033785m | *Manes.01G215000.1* | *MeGRX785* | MeGRXS1-related | Chromosome01:30421882..30422775 |

**Fig. 2.**
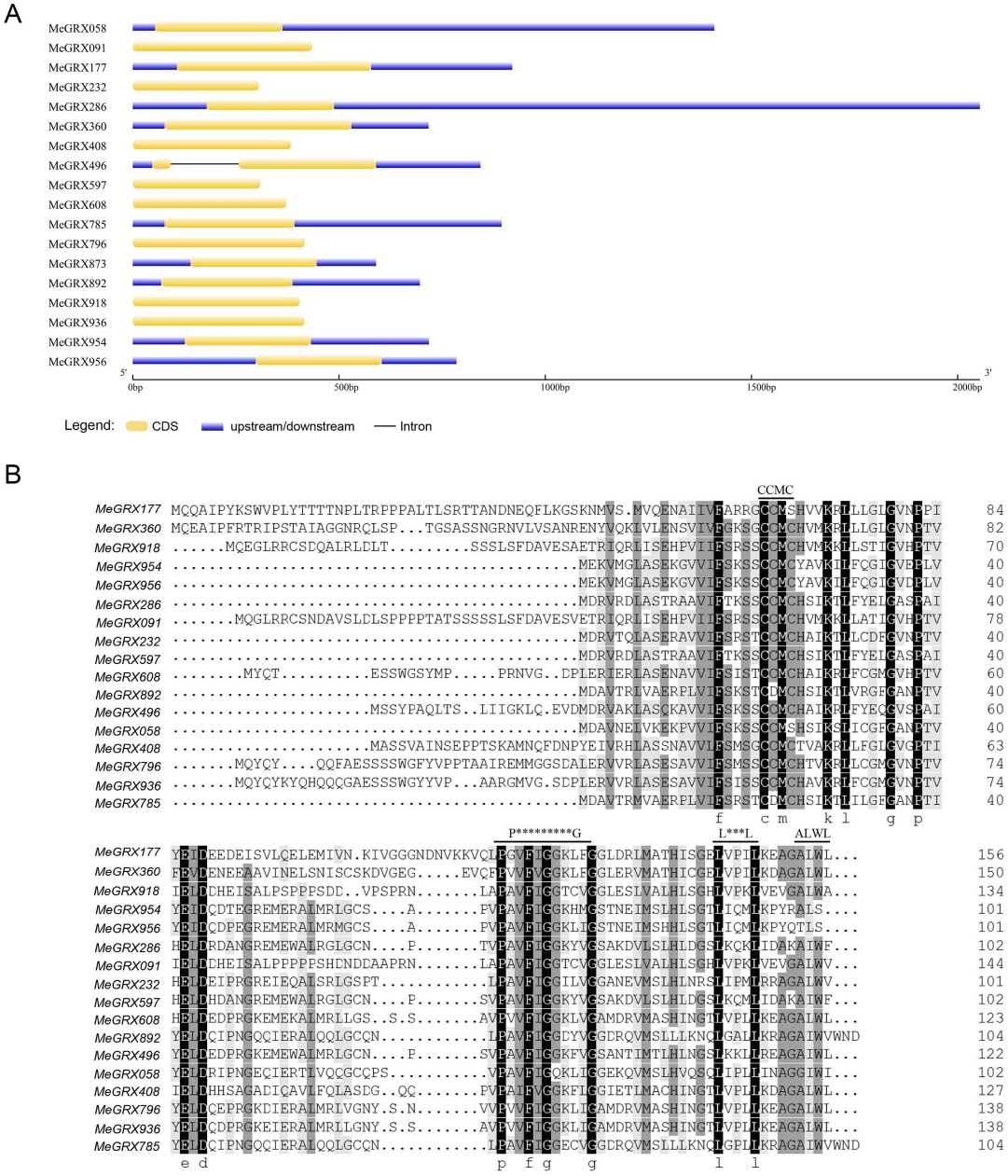
Gene structure and protein sequences alignment of cassava CC-type GRXs. A. The exon-intron structure of cassava CC-type GRXs. Analysis was carried out with GSDS (http://gsds.cbi.pku.edu.cn); B. Protein sequences alignment of cassava CC-type GRXs. The editing of aligned sequences among cassava CC-type GRXs was performed using AlignX (Vector NTI suite 10.3 Invitrogen). Black boxes indicate conserved indentify positions, gray box indicate similar positions. The letters above the sequence indicate motif name.

### Identification of drought- and ABA-inducible CC-type GRX genes in cassava cultivars

As a tropical crop, cassava is well adapted to interment drought, but hypersensitive to cold (An *et al*. 2012; Okogbenin *et al*. 2013; Xia *et al*. 2014; Zeng *et al*. 2014; Zhao *et al*. 2014). Previous studies used high quality RNA-seq and iTRAQ-based proteomic datasets to examine genes responsive to drought resistance in cassava (W.Hu *et al*. 2015b; W.Hu *et al*. 2015a; Xia *et al*. 2014; Zeng *et al*. 2014; Zhao *et al*. 2014). To investigate the role of CC-type GRXs in response to drought in cassava, we used eight arrays of our previously reported transcriptomic data (Wei Hu *et al*. 2016). RNA-seq datasets are available at NCBI and the accession numbers are listed in Table S4. Hierarchical expression clustering (FPKM) showed that the CC-type GRX expression patterns in response to drought in cassava cultivars Arg7 and SC124 grouped in three clusters (Fig.3A). Cluster III included CC-type GRXs induced by drought only in leaves. To detail the expression of CC-type GRXs in the drought response in cassava leaves, we performed a qPCR analysis to investigate the expression changes of genes in cluster III under drought and re-water treatments. For this analysis, we selected six drought-inducible CC-type GRXs (*MeGRX058, 232, 360, 496, 785*, and *892*) from cluster III on the basis of RNA-seq. We collected leaves from plants of two cassava cultivars under drought stress for eight (D8) or 14 days (D14), and D14 plants re-water 24 hours later (RW). We used leaves from well-water cassava plants as control (DC). As expected, drought stress up-regulated the expression of all six CC-type *GRXs* in both Arg7 and SC124 leaves (Fig. 3B, C). It indicates that CC-type GRXs may play conserved roles in drought response of cassava different cultivars. The expression of *MeGRX360, MeGRX785*, and *MeGRX892* was the highest in D14 plants. In contrast, the expression of *MeGRX058* and *MeGRX232* was the highest in D8 plants. These results indicated that CC-type GRXs may regulate drought response through different mechanisms in cassava.

**Fig. 3.**
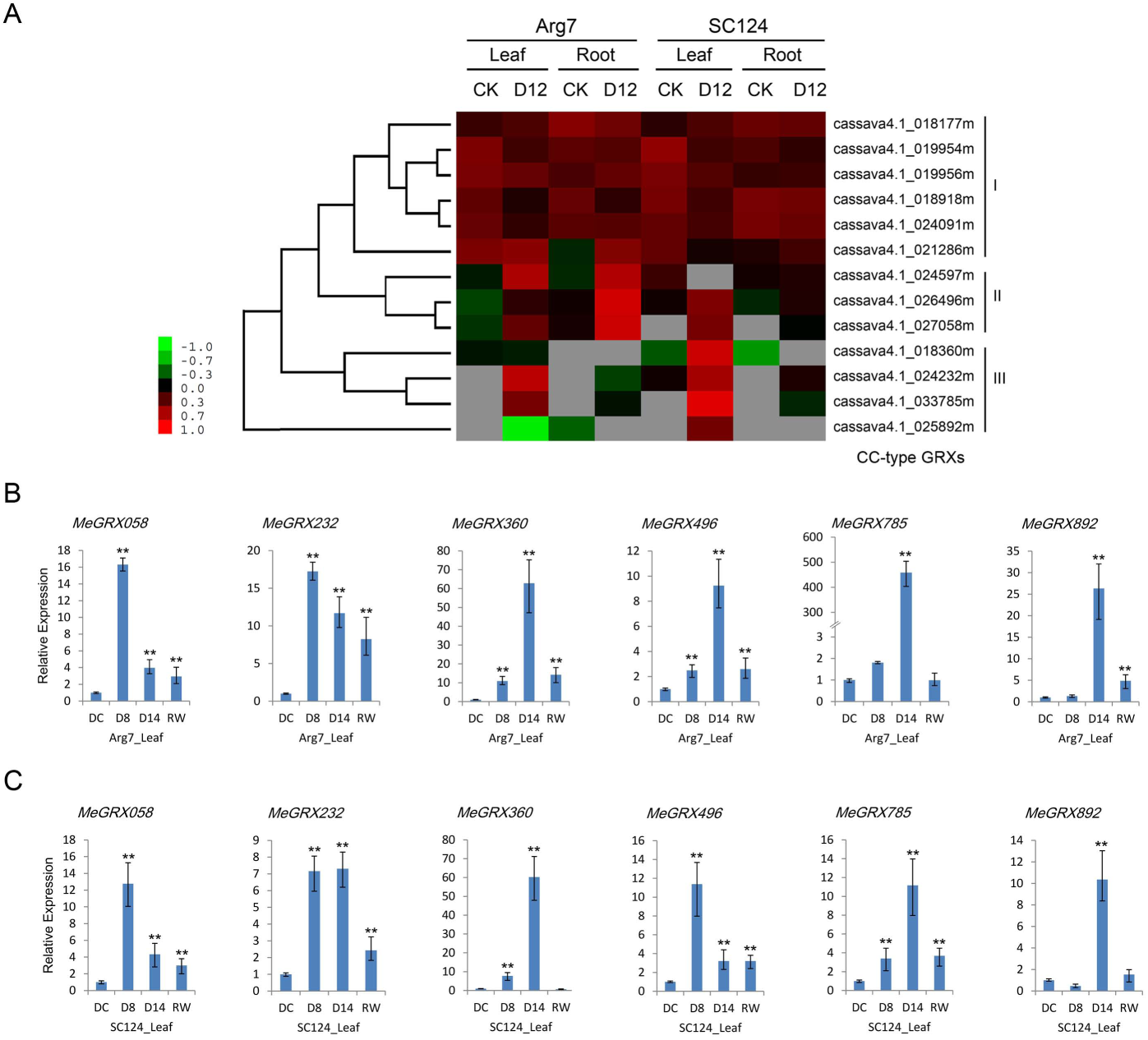
Figure 3 Expression analysis of CC-type GRXs in cassava Arg7 and SC124. (a) Heat map represent expression of CC-type GRXs in Arg7 and SC124. Hierarchical clustering was performed using Cluster3.0 based on the RNA-seq data from drought stressed Arg7 and SC124 plants. Heat map was built using TreeView1.0.4. (b) qPCR analysis of six drought-inducible CC-type GRXs in drought stressed fifth leaves of Arg7. (c) qPCR analysis of six drought-inducible CC-type GRX genes in drought stressed fifth leaves of SC124. DC: control; D8: 8 day after the start of the drought stress treatment; D14: 14 day after the start of the drought stress treatment; RW: 1 day after re-watering at the ending of drought treatment. Expression levels of the six CC-type GRXs were normalized against DC. Biological triplicates were averaged and significance of differences between treatments and control were analyzed using the Student’s t-test (**, p≤ 0.01). Bars represent the mean ± standard error.

The phytohormone ABA regulates many important processes in plants, especially in relation to environmental stress responses (Nakashima and Yamaguchi-Shinozaki 2013; Sharp and LeNoble 2002; Wilkinson and Davies 2002). Numerous drought-responsive genes have been described as ABA-inducible (Nakashima and Yamaguchi-Shinozaki 2013). A conserved cis-element, designated the ABA-responsive element (ABRE), is present in the promoter region of most ABA-inducible genes (Fujita *et al*. 2011). Analyzing the 1.5 kb up-stream region of our six drought responsive CC-type *GRXs*, we found ABREs in the promoter region of *MeGRX232, MeGRX360, MeGRX496, MeGRX785*, and *MeGRX892*. Thus, we inferred that these genes may respond to ABA. We performed qPCR analysis to confirm this hypothesis. Interestingly, ABA application up-regulated the expression of these six CC-type GRXs in both Arg7 and SC124 leaves (Fig. 4). The data suggest these CC-type *GRXs* may play roles in ABA signal transduction pathways in cassava.

**Fig. 4.**
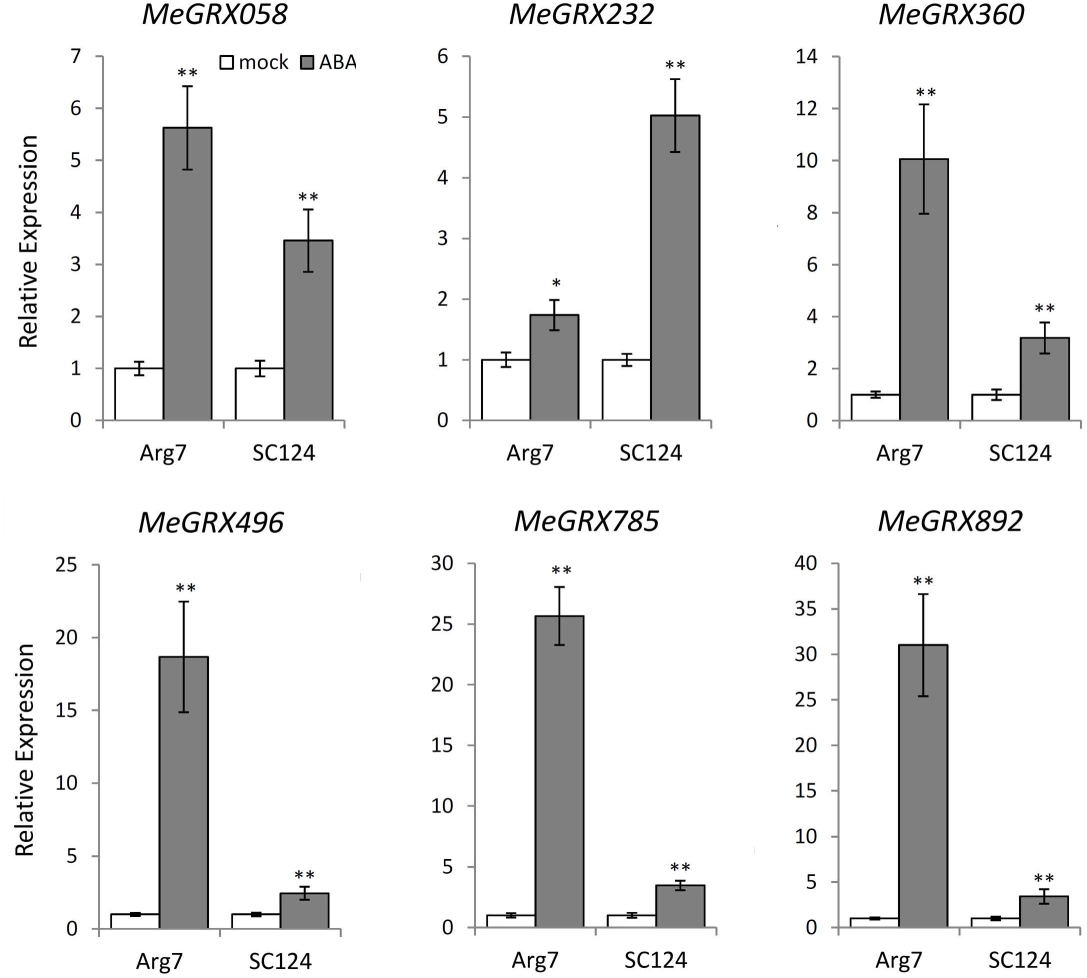
Expression analysis of cassava CC-type GRXs respond to exogenous ABA in leaves. The expression level of these six CC-type GRX genes were set to 1 in DC. Biological triplicates were averaged and significance of differences between treatments and control were analyzed using the Student’s t-test (**, p ≤ 0.01; *, 0.01 < p ≤ 0.05). Bars represent the mean ± standard error.

### Overexpression of *MeGRX232* in *Arabidopsis* confers seedling development insensitive to ABA and affects root architecture under mannitol stress

Most *Arabidopsis* CC-type GRX proteins localize in the cytosol or in the nucleus (Couturier *et al*. 2011; Rouhier *et al*. 2007; Xing *et al*. 2005; Xing and Zachgo 2008). The sub-cellular localization of these proteins is essential for their function (Li *et al*. 2009). We tagged the cDNA of our six drought-responsive CC-type GRXs with GFP at the C-terminus to analyze their cellular localization (Fig. S1). We imaged our *MeGRX:GFP* fusions in transiently transformed *Nicothiana benthamiana* leaf epidermis, detecting fluorescence in both the cytosol and the nucleus (Fig. S1). In *Arabidopsis*, the nuclear localization of ROXY1 is required for its function in petal development (Li *et al*. 2009). Thus, the sub-cellular indicates that these drought-inducible CC-type GRXs may function in the nucleus during stress responses.

To investigate the functions of drought-inducible MeGRXs in plant, we heterologous overexpressing four of these genes that is *MeGRX058, 232, 360*, and *785* in *Arabidopsis* (Fig. S2). Overexpression of *MeGRX785* caused ABA and mannitol susceptibility of seed germination in *Arabidopsis*, while in contrast, *MeGRX232-OE* plants was insensitive to ABA and mannitol (Fig. S2). Three independent lines of *MeGRX232-OE* transgenic *Arabidopsis* have been used in ABA and osmotic stress analyses. And the transgenic *Arabidopsis* that contained vector (*pG1300*) were used as control. We found that ABA did not affect the seed germination and seedling development of *MeGRX232-OE* transgenic plants (Fig. S3). We infer that overexpression of *MeGRX232* may cause ABA insensitivity in *Arabidopsis*. Next, 5-day-old seedlings of *MeGRX232-OE* transgenic *Arabidopsis* were grown on MS medium supplement with 0μM ABA (mock) or 5μM ABA, respectively. After 10 days grown on mock medium, no visible phenotypic differences between *MeGRX232-OE* and vector plants were observed (Fig.5A). On ABA-supplement medium, the growth of vector plants was significantly inhibited, while the growth of *MeGRX232-OE* plants were less inhibited (Fig.5A). The rosette diameter of *MeGRX232-OE* plants was ∼50% higher than that of vector plants (Fig.5B). Also, the primary root of *MeGRX232-OE* plants was ∼48% longer than that of vector plants (Fig.5C). Our data address the issue that overexpression of *MeGRX232* caused ABA insensitivity in transgenic *Arabidopsis*.

**Figure 5.**
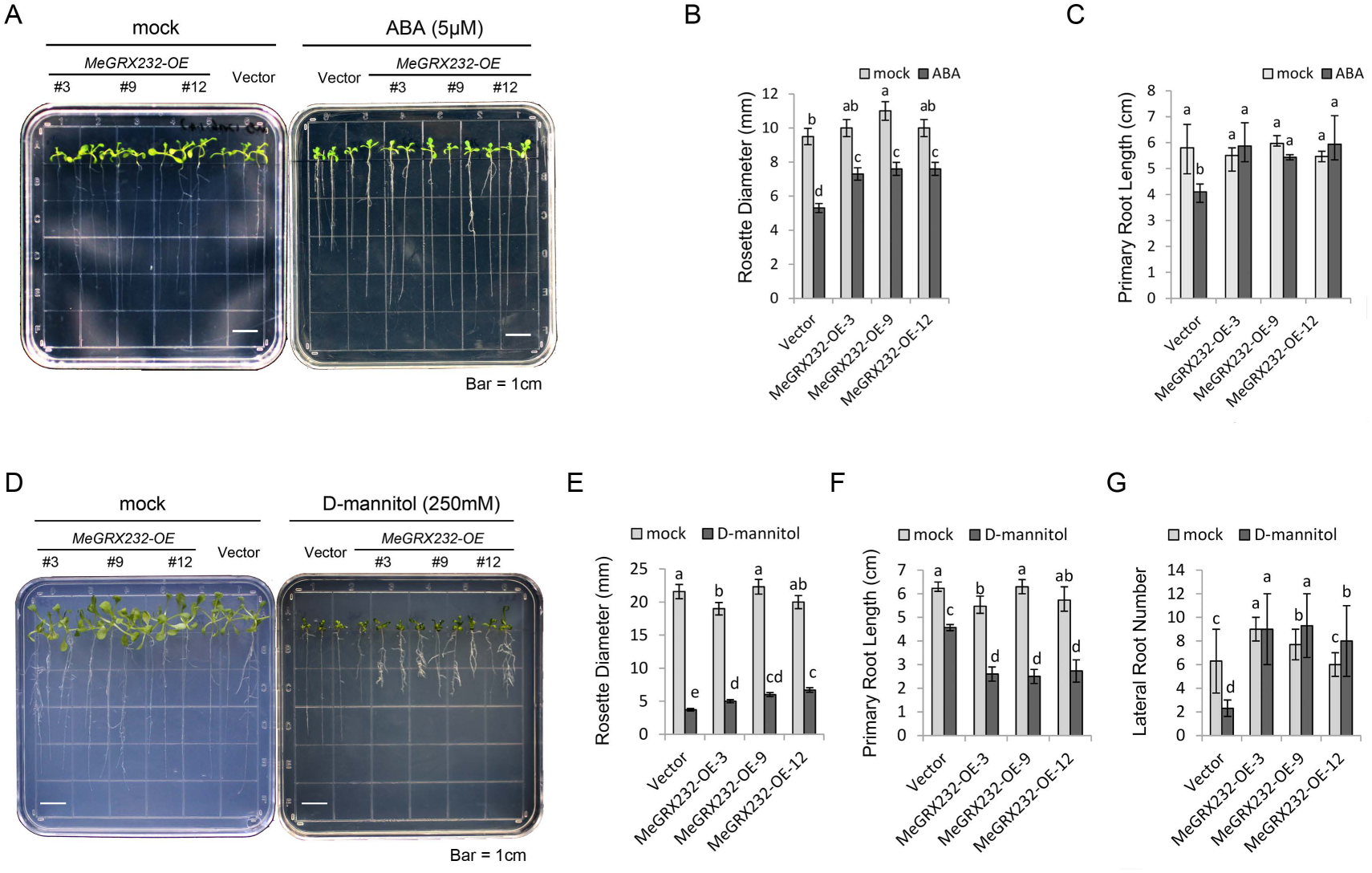
Effects of ABA and mannitol on seedling development of *MeGRX232-OE Arabidopsis*. (a) Post-germinated seedlings development of transgenic plants on MS medium supplemented with 0 (mock) and 5μM ABA, respectively. The plants that contained pG1300 (Vector) were used as control. (b), (c) Rosette diameter and primary root length of transgenic *Arabidopsis* in (a). (d) Post-germinated seedlings development of transgenic plants on MS medium supplemented with 0 (mock) and 250mM D-mannitol, respectively. The plants that contained pG1300 (Vector) were used as control. (e), (f), (g) Rosette diameter, primary root length, and lateral root number of transgenic *Arabidopsis* in (d). Biological triplicates were averaged and significance of difference between treatments and control was analyzed using the Duncan’s multiple range tests. Error bars show standard errors for three independent replicates. Different letters represent a significant difference at p <0.05.

To analysis mannitol tolerance of transgenic *Arabidopsis*, 5-day-old seedlings of transgenic plants were grown on MS medium supplement with 0 mM (mock) and 250 mM D-mannitol. After 15 days, the *MeGRX232-OE* and vector plants show no significant difference on mock medium (Fig.5D). While root system of *MeGRX232-OE* plants were dramatically affected by 250 mM D-mannitol (Fig.5D). The rosette diameter of *MeGRX232-OE* plants was higher than that of vector plants, ranged from ∼25% to ∼50% (Fig.5E). However, the primary root elongation in *MeGRX232-OE* plants decreased ∼36% compared with that in vector plants (Fig.5F). Under mannitol stress, *MeGRX232-OE* plants have more lateral roots than vector plants. We found that the lateral root number increased ∼4 fold in *MeGRX232-OE* plants, compared to vector plants (Fig.5G). Collectively, the data indicate that overexpression of *MeGRX232* affected root architecture in *Arabidopsis* under mannitol stress.

### *MeGRX232* confers drought hypersensitivity in soil-grown plants via impairing ABA-dependent stomatal closing

To further investigate the role of *MeGRX232* in drought tolerance, *MeGRX232-OE* and vector plants grown in soil were used. The vector and three independent lines of *MeGRX232-OE* plants were grew in one pot under normal conditions. 21-day-old plants in soil have been treated by water withholding (Fig.6A). When exposed to water deficit for 21 days, all treated plants displayed severe wilting (Fig.6B). After drought treatment, plants were re-watered and cultured for five more days. The *MeGRX232-OE* lines displayed a significantly lower survival rate than vector plants (Fig.6C). It indicates that overexpression of *MeGRX232* caused drought hypersensitivity in *Arabidopsis* under soil culture conditions. This might result from a more rapid loss of water in *MeGRX232-OE* plants than in vector plants (Fig.6D).

**Figure 6.**
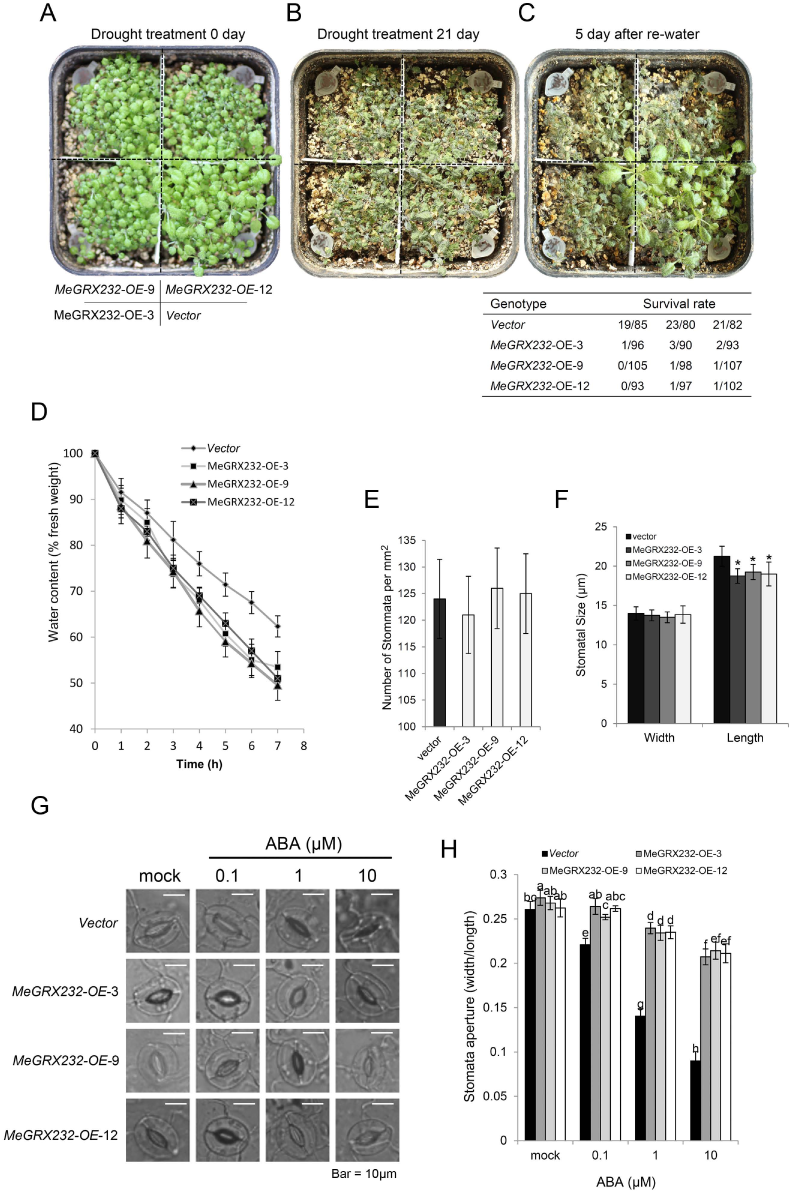
Drought tolerance analyses of transgenic *Arabidopsis* grown in soil and effect of ABA on transgenic *Arabidopsis* with respect to stomatal aperture. (a), (b), (c) Drought responses of transgenic plants. Survival rate has been calculated from three independent experiments. (d) Water loss rate of transgenic plants. Biological triplicates were averaged. Error bars show standard errors for three independent replicates. (e) Stomatal distribution in leaves of transgenic *Arabidopsis*. (f) Stomatal size of transgenic *Arabidopsis*. (g) Effects of ABA on stomata of abaxial leaf epidermal peels were observed. The abaxial leaves were treated by ABA with different concentrations, respectively. (h) Stomatal aperture measurement after ABA treatment in (g) was carried out by recording width to length ration. Biological triplicates were averaged and significance of difference between treatments and control was analyzed using the Duncan’s multiple range tests. Error bars show standard errors for three independent replicates. Different letters represent a significant difference at p <0.05.

For water loss mainly depends on stomatal regulation, we infer that *MeGRX232* affect stomatal density or movement therefore increase water loss in transgenic *Arabidopsis*. However, there are no obviously stomatal density differences between vector and *MeGRX232-OE* plants (Fig.6E). But the stomata apertures (width/length) of MeGRX232-OE plants are higher than that of vector plants (Fig.6F), suggesting *MeGRX232* may affect stomatal closure during drought stress. Thus, we performed ABA-induced stomatal closing assays to test our hypothesis. Without ABA (mock), the *MeGRX232-OE* lines showed similar stomatal aperture as vector plants (Fig.6G, H). This indicates that *MeGRX232* overexpressing did not affect stomatal aperture in *Arabidopsis* under normal conditions. When treated with 0.1 μM ABA, the stomata of vector plants displayed a significant closing, whereas the stomata of *MeGRX232-OE* lines open widely as that in mock (Fig.6G, H). Furthermore, with increasing concentrations of ABA to 1 μM and 10 μM, the stomatal aperture of vector plant exhibited dramatically reducing. On the contrary, the *MeGRX232-OE* lines performed relatively less closing of stomata (Fig.6G, H). It suggests that overexpression of *MeGRX232* resulted impairment of ABA-dependent stomatal closing. Therefore, the more rapid water loss observed in *MeGRX232-OE* lines should mainly be ascribed to impaired stomatal closure.

### Overexpression of MeGRX232 caused more ROS accumulation in guard cells

CC-type GRX GRXS13 is critical for ROS production during photooxidative stress (Laporte *et al*. 2012). To determine whether MeGRX232 affect cell redox homeostasis, we measured the MDA content in stress transgenic plants. We found a higher MDA content in *MeGRX232-OE* plants than in vector plants (Fig.7A). Furthermore, we found more ROS signals have been stained by DAB in *MeGRX232-OE* plants under normal conditions (Fig.7B). Together, these data indicate the possibility that overexpression of *MeGRX232* caused more ROS accumulation in transgenic *Arabidopsis* leaves.

**Figure 7.**
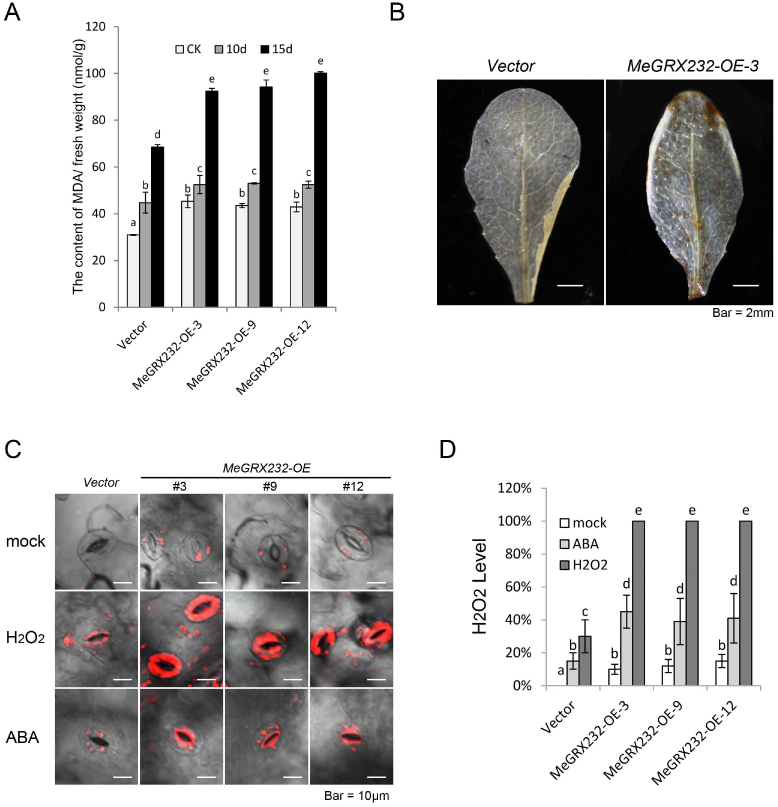
ROS accumulation analysis in transgenic *Arabidopsis*. (a) MDA content of transgenic *Arabidopsis* during drought treatment. (b) DAB staining of transgenic *Arabidopsis* leaves. (c) ROS accumulation in guard cells of leaves from transgenic *Arabidopsis*. (d) Quantification of ROS levels in guard cells of transgenic *Arabidopsis*. The fluorescent intensity in *MeGRX232-OE Arabidopsis* after H_2_O_2_ treatment was taken as 100%. More than 30 stomata of each leaf were calculated and significance of difference between treatments and control was analyzed using the Duncan’s multiple range tests. Error bars show standard errors for three independent replicates. Different letters represent a significant difference at p <0.05.

As H_2_O_2_ promotes leaf stomatal closure acting downstream of ABA (Pei *et al*. 2000). H_2_O_2_ accumulation in guard cells was measured by a fluorescence dye, 2’, 7‘-dichlorodihydrofluorescein diacetate (DCFH-DA) under exogenous ABA and H_2_O_2_ treatment. The *MeGRX232-OE* guard cells show obvious H_2_O_2_ accumulation, in contrast, no H_2_O_2_ accumulation in vector plants guard cells have been detected (Fig.7C, D). After treated by H_2_O_2_ for three hours, both *MeGRX232-OE* and vector plant guard cells show significant H_2_O_2_ accumulation, but the H_2_O_2_ levels in *MeGRX232-OE* guard cells are much higher than that in vector plant guard cells (Fig.7C, D). After treated for three hours, more H_2_O_2_ have been induced by ABA in *MeGRX232-OE* guard cells compared to vector plant guard cells, and the H_2_O_2_ have been accumulated in membrane of *MeGRX232-OE* guard cells (Fig.7C, D). Thus, we can conclude that overexpression of *MeGRX232* in *Arabidopsis* caused more ROS production in guard cells.

### MeGRX232 is interacts with TGA5 and MeTGA074

Most CC-type GRXs play roles in organ development and plant defense via interaction with TGA factors (Hong *et al*. 2012; Li *et al*. 2009; Li *et al*. 2011; Zander *et al*. 2012). TGA factors regulate genes that involved in both biotic and abiotic stress (Sham *et al*. 2014). It is necessary to identify the interactors of MeGRX232 in *Arabidopsis* and cassava. We fused MeGRX232 with the GAL4 DNA-binding domain (BD) sequence in *pGBKT7* (Clontech) and then transformed the result construct into yeast strain Y187. The *pGBKT7* vector was used as negative control. However, yeast cells harboring *MeGRX232:pGBKT7* activated X-α-gal on SD/-Trp / X-α-gal medium (Fig.8A), suggesting MeGRX232 has transcriptional activation ability. CC-type GRXs need to interact with GSH to catalyze essential biosynthesis reactions by its redox regulation (Xing and Zachgo 2008). Therefore we created a MeGRX232 mutant by replacing GSH binding site. As is shown in Fig. 8c, the MeGRX232 mutant did not activated X-α-gal on the medium. It suggests that the GSH binding site is required for transcriptional activation ability of MeGRX232. A possible explanation is that MeGRX232 could binding and modify transcription factor depending on GSH in yeast.

**Figure 8.**
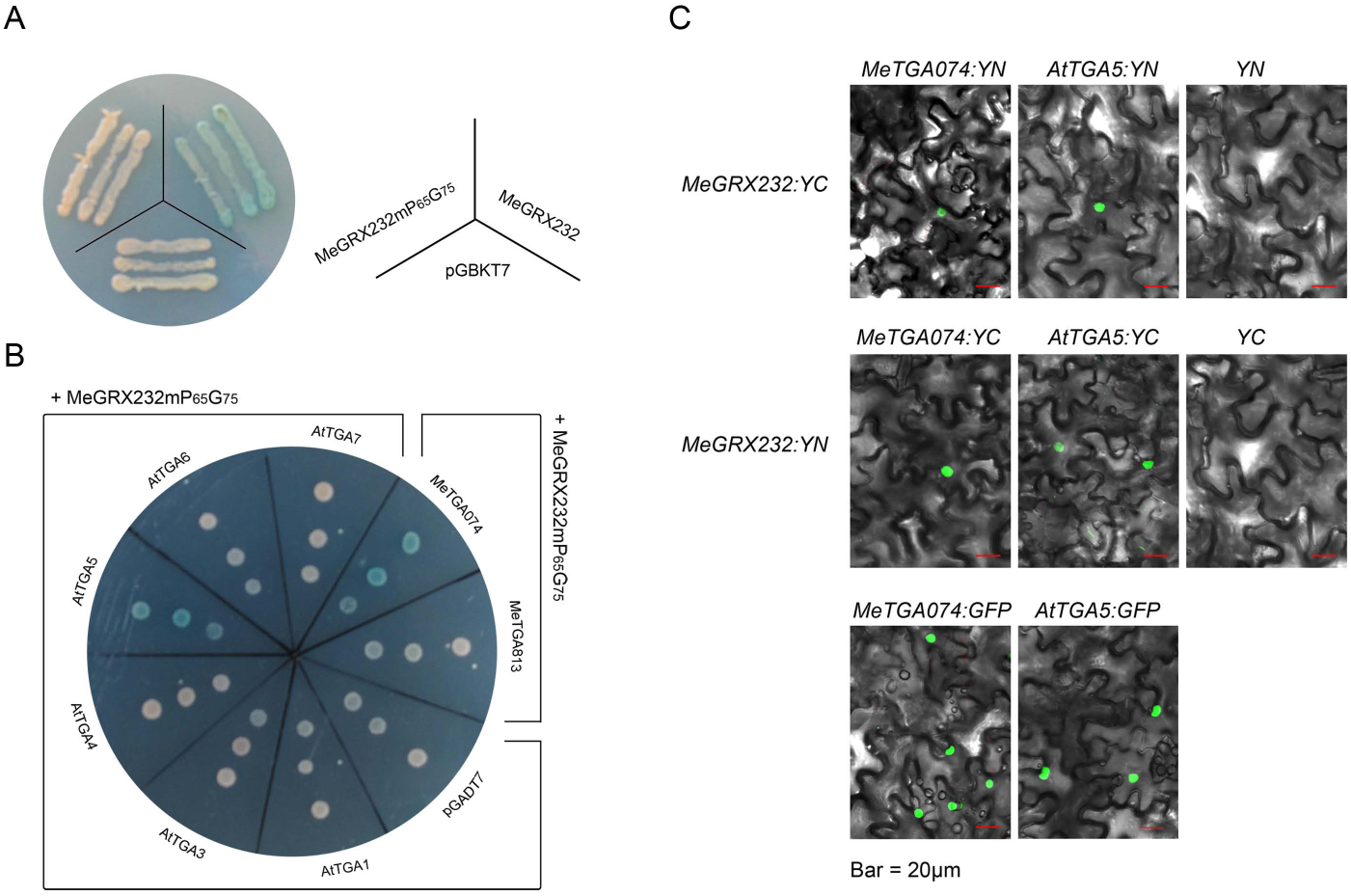
Identification of MeGRX232 interactors in *Arabidopsis* and cassava. (c) Autonomous transactivation analysis of MeGRX232 in yeast. MeGRX232mP6_5_G6_5_ indicate mutant in MeGRX232 GSH binding site. (d) Analysis of interaction between MeGRX232 mP_65_G_65_ and TGA factors by yeast two hybrid system. (e) BiFC analysis of the interactions between MeGRX232 and TGAs identified by (d) in transiently transformed *N. benthamiana* leaves. Green fluorescence in nucleus was detected for interactions of MeGRX232 with MeTGA074 and AtTGA2, respectively. As a negative control, co-expression of MeGRX232:YN with free YC, and MeGRX232:YC with free YN failed to reconstitute a fluorescent YFP chromophore. Expression of MeTGA074:GFP and AtTGA2:GFP in transiently transformed *N. benthamiana* as positive controls.

Subsequently, six TGA factors including TGA1, TGA3, TGA4, TGA5, TGA6, TGA7 in *Arabidopsis* and two TGA factors (MeTGA074 and MeTGA813) in cassava have been fused with GAL4 activation domain (AD) sequence in pGADT7 (Clontech). The resulted AD:TGA constructs and BD:MeGRX232mP_65_G_75_ were pairwise co-transformed into yeast Y187, respectively. Yeast cells that harboring both *AD:TGA* and *BD:MeGRX232mP_65_G_75_* pair plasmids were grown on SD/ -Trp/ -Leu/ X-α-gal medium (Fig.8B). The yeast cells containing pairwise plasmids AD:TGA5/BD:MeGRX232mP_65_G_75_ and AD:MeTGA074/BD:MeGRX232mP_65_G_75_ activated X-a-gal. It suggests that MeGRX232 could respectively interact with TGA5 or MeTGA074.

To further investigate the interactions between MeGRX232 and TGA5/MeTGA074 *in planta*, we employed BiFC. Nuclear fluorescence co-expression of *MeGRX232* and *TGA5/MeTGA074* has been detected in epidermal cells (Fig.8C). The *in planta* nuclear interactions of MeGRX232 with TGA5/MeTGA074 suggest that this CC-type GRX might functions in *Arabidopsis* and cassava by nuclear interacting with TGA5/MeTGA074. We created a phylogenetic tree based on TGA protein sequences in *Arabidopsis* and cassava (Fig. S4). We found that MeTGA074 is a member of clade II TGA, closely to TGA5. Together, our data suggest that *MeGRX232* may regulate drought response via interaction with TGA5/MeTGA074.

### MeGRX232 regulates a group of genes involved in stress and redox homeostasis in *Arabidopsis*

To understand the effects of the *MeGRX232* overexpressing on gene expression in *Arabidopsis*, a microarray analysis has been performed using Affymetrix Arabidopsis ATH1 Genome Array. Three independent lines of *MeGRX232-OE* and vector *Arabidopsis* grew in soil under normal conditions were used. We found that transcription levels of 2674 genes were altered significantly (with more than a twofold change; P value < 0.05) in *MeGRX232-OE* lines compared with vector lines under normal conditions (Table S5). 1264 genes were up-regulated, whereas 1410 genes were down-regulated. The relative expression levels of these genes were shown by the heat map (Fig.9A). Gene ontology (GO) analysis results showed that many stress responsive genes have been affected by *MeGRX232-OE Arabidopsis* (Fig.9B). 27 more abundant GO categories (q-value < 10^-5^) including categories that response to abiotic, biotic stress, and phytohormone stimulus in *MeGRX232-OE Arabidopsis* were exhibited here. Interestingly, nearly two hundred transcription factors have been affected by *MeGRX232-OE Arabidopsis* (Fig.9B). We found that 192 oxidative stress-related, 44 drought related, and 53 ABA related genes were significantly altered in *MeGRX232-OE* plants. Nevertheless, there was three genes overlap among the genes in response to drought, oxidative stress and ABA (Fig.9C), indicating that a specific regulatory mechanism which dependent on ABA-ROS crosstalk conferred by MeGRX232 is presented in response to drought. Moreover, there was three drought related and three oxidative stress related genes were overlapped to the genes that involved in JA/ET signal transduction respectively (Fig.9C), suggesting the MeGRX232 may play roles in drought response depending on regulation of JA/ET pathway.

**Figure 9.**
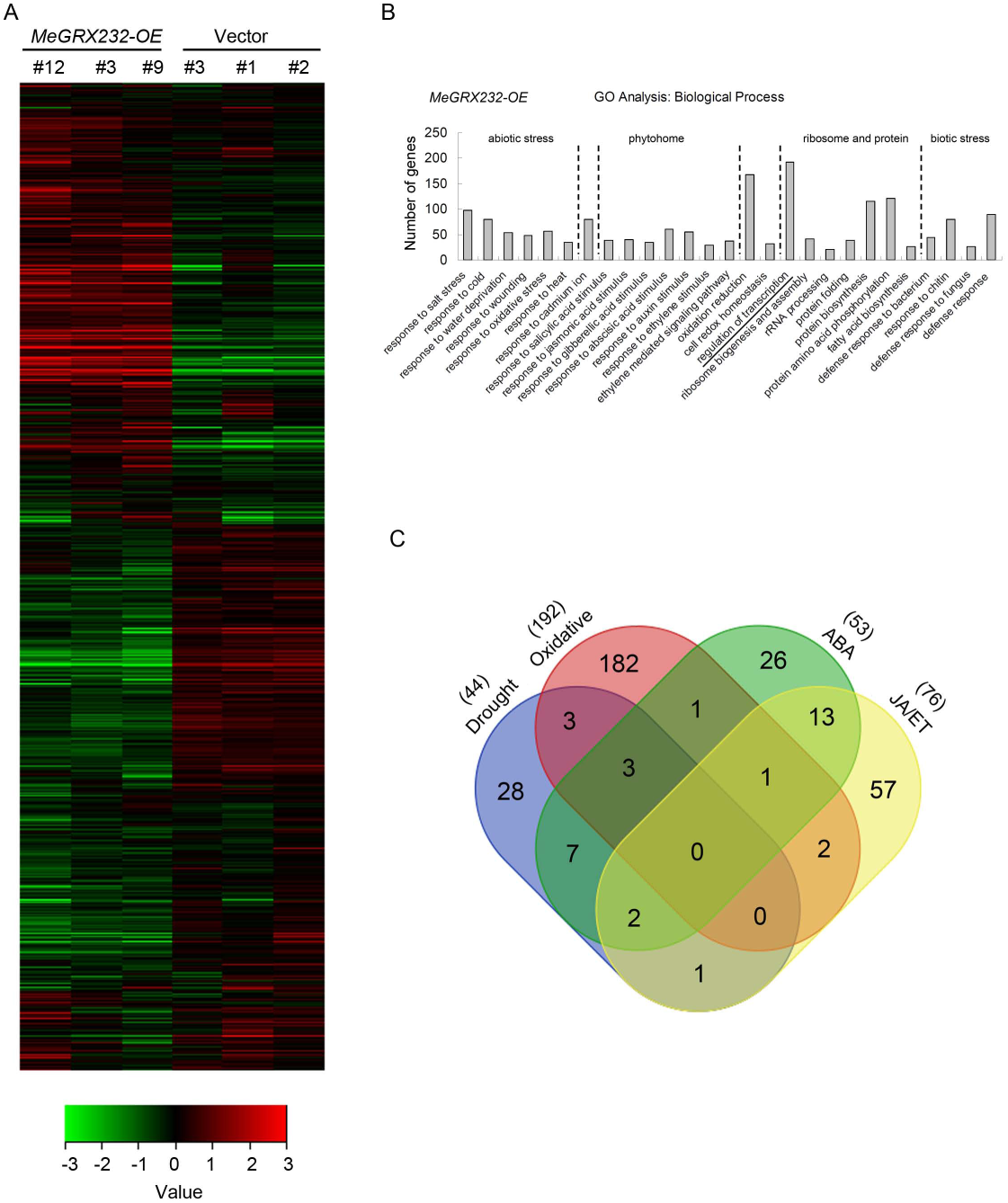
Gene expression profiles in *MeGRX232-OE* transgenic *Arabidopsis* and role of MeGRX232 regulators in drought response. (a) Heat map represent gene expression between *MeGRX232-OE* and control (Vector) plants. The data was processed and normalized as described in Materials and Methods. Hierarchical clustering of significantly expressed genes is displayed by average linkage. The figure was drawn by TreeView software. (b) GO analysis of *MeGRX232-OE* induced genes in *Arabidopsis*. Comparison of GO terms identified from the differentially expression genes identified in SAM analysis. GO tags were selected according to the significance (p-value <10^-5^). Numbers on y-axis indicate gene numbers of the GO tag. (c) Venn diagram showing the overlap between MeGRX232-OE regulated genes in response to different stress and signals.

## Discussion

CC-type GRXs is a land plant specific subgroup of GRX family, derived from the CPYC subgroup (Ziemann *et al*. 2009). Phylogenetic data (Fig. 1) showed that cassava CC-type GRXs developed from two CPYC members (cassava4.1_018918m and cassava4.1_024091m). During land plant evolution, CC-type GRXs gained new functions (Rouhier *et al*. 2006; Wang *et al*. 2009; Ziemann *et al*. 2009). In *Arabidopsis*, CC-type GRXs play an important role in petal development and biotic stress responses (Ndamukong *et al*. 2007; Xing *et al*. 2005). CC-type GRXs function in the regulation of ethylene responsive genes through the interaction with TGA factors (Garg *et al*. 2010; La Camera *et al*. 2011; Meyer *et al*. 2012; Ndamukong *et al*. 2007; Wang *et al*. 2009; Zander *et al*. 2012). The interaction between these proteins and TGA factors depends on their C-terminal L**LL and ALWL motifs (Li *et al*. 2009; Li *et al*. 2011; Zander *et al*. 2012). These motifs were also present in almost all cassava CC-type GRXs except MeGRX954 and MeGRX956 (Fig. 2).

To date, no CC-type GRX has been identified as a regulator of drought response in cassava. Based on our previously reported RNA-seq data (Wei Hu *et al*. 2016), we identified six drought stress inducible CC-type GRXs in leaves of cassava using qPCR analysis (Fig. 3). Under drought stress, ABA concentrations increases and, in turn, induces gene expression (Huang *et al*. 2011). In our study, ABA stress up-regulated the expression of these six drought-inducible CC-type GRX genes in leaves of both Arg7 and SC124 plants (Fig. 4). Thus, we believe that CC-type GRXs might be playing roles in cassava drought response in an ABA-dependent manner. Therefore, understanding the molecular mechanisms controlled by CC-type GRXs may provide an effective method for genetic improvement of drought stress resistance for cassava and other crops. We over-expressed four drought-inducible CC-type GRX genes (*MeGRX058, MeGRX232, MeGRX360* and *MeGRX785*) in *Arabidopsis* under control of the *CMV 35S* promoter (Fig. S1). Indeed, when we compared the seed germination of and *MeGRX232-OE, MeGRX785-OE* and wild type *Arabidopsis* on MS media supplemental with 2μM ABA, significant inhibition of seed germination was observed in *MeGRX785-OE* only (Fig. S1), and the seedling development of *MeGRX232-OE Arabidopsis* was insensitive to ABA and mannitol (Fig. 5), indicating that members of drought-inducible CC-type GRXs may play different roles in responses to drought in cassava.

Recently, several abiotic stress related CC-type GRXs have been identified in *Arabidopsis* and rice (El-Kereamy *et al*. 2015; Gutsche *et al*. 2015; La Camera *et al*. 2011; Laporte *et al*. 2012; R.Sharma *et al*. 2013). A splicing variant of *AtGRXS13* (*ROXY19*) involved in the protection against photo oxidative stress in *Arabidopsis* (Laporte *et al*. 2012). Overexpression of the rice CC-type GRX *OsGRX8*, which is generally induced by auxin and abiotic stress enhances tolerance to ABA and abiotic stresses in *Arabidopsis* (R.Sharma *et al*. 2013). Here, the overexpression of *MeGRX232* in *Arabidopsis* caused tolerance to ABA and mannitol on the seal agar plates (Fig. 5). However, the *MeGRX232-OE Arabidopsis* showed drought hypersensitivity in soil-grown condition (Fig. 6). Then we found the drought hypersensitivity is partly resulted by impairment of ABA-dependent stomatal closure in *MeGRX232-OE Arabidopsis* (Fig.6G). Therefore, the inverse phenotypes of *MeGRX232-OE Arabidopsis* were possibly ascribed to the different stress conditions: in sealed agar plates, the transpiration of seedlings is almost negligible (Verslues *et al*. 2006), whereas in open environment, the *MeGRX232-OE* plants show higher water loss rate (Fig.6D).

Overexpression of *MeGRX232* resulted in hypersensitivity to drought and caused a higher water loss rate, which led us to suppose that MeGRX232 is involved in stomatal movement. ABA-induced stomatal closing was impaired in *MeGRX232-OE* plants (Fig. 6), suggesting that MeGRX232 plays a role in the inhibition of stomatal closing. But the main function of MeGRX232 in stomatal regulation seems not to only inhibit stomatal closing, because it is contradictory for cassava to induce the expression of *MeGRX232* under dehydration conditions to inhibit stomatal closing. During drought treatment, the high ROS accumulation is essential for abscission zone initial in cassava petiole (Liao *et al*. 2016a). *MeGRX232* was induced by drought not only in leaves but also in abscission zone (Fig. S5), and overexpression of *MeGRX232* caused more ROS accumulation in *Arabidopsis* (Fig. 7), suggesting a potentially role of *MeGRX232* in ROS accumulation during abscission zone formation in cassava. It will be of interesting to investigate whether this gene is involved in leaf abscission in cassava.

The interaction with TGA factors is necessary for CC-type GRX functions in plants (Li *et al*. 2009; Li *et al*. 2011; Zander *et al*. 2012). In *Arabidopsis*, TGA factors have been classified to five subgroups, clade I, II, III, IV, and V. TGA2, 5, 6 are members of clade II TGAs, which are essential activators of jasmonic acid/ethylene-induce defense responses (Kesarwani *et al*. 2007; Stotz *et al*. 2013; Zander *et al*. 2010; Zander *et al*. 2012) and act as a key regulator role in plant responses of abiotic stresses such as drought, cold, and oxidative stress (Sham *et al*. 2014). *Arabidopsis* CC-type GRX GRX480/ROXY19 could interact with TGA2, 5, 6 (Zander *et al*. 2012). TGA2 could interact with GRXS13, and act as repressors of GRXS13 expression in response to biotic stress (La Camera *et al*. 2011). Here, we found that MeGRX232 nuclear interacted with TGA5 in *Arabidopsis* and MeTGA074 in cassava respectively (Fig. 8). In *Arabidopsis*, GRX480 regulated the expression of ERF (Ethylene Response Factor) factors through interaction with TGA2/5/6 (Ndamukong *et al*. 2007; Zander *et al*. 2012). We found that nearly two hundred transcription factors including ERF factors had been affected by MeGRX232-OE in *Arabidopsis* (Fig.9B, Table S5). A nuclear export signal (NES) should be tag to MeGRX232 to eliminate its nuclear localization to investigate whether the MeGRX232 regulated ERFs through nuclear interaction with TGA5 in *Arabidopsis*. The presence of TGA binding elements (TGACG) in the sequence of MeGRX232 promoter suggests that members of TGA factors could directly bind to the promoter (Table S6). It will be of interest to further study the mechanism by which MeGRX232 respond to drought via interaction with MeTGA074 in cassava.

In summary, we demonstrate that *MeGRX232* plays a key role in regulating stomatal closure. The expression of *MeGRX232* in cassava leaf can be induced by drought and ABA. Overexpression of *MeGRX232* results increased ROS accumulation in guard cells, and confers drought hypersensitivity by inhibition of ABA-dependent stomatal closing in *Arabidopsis*. As a CC-type GRX, MeGRX232 could interact with *Arabidopsis* TGA2 and cassava MeTGA074 factors, and regulated the expression of ABA and oxidative stress related genes in *Arabidopsis*. Our study demonstrates that CC-type GRXs may functions in ABA-ROS signal transduction in drought response of cassava. It will contribute to an enhanced understanding of the specific mechanisms that elucidate the roles of CC-type GRXs involved in drought response in cassava.

## Supplementary data

Supplementary figure S1. Protein localization analysis of six drought-responsive CC-type GRXs.

Supplementary figure S2. Identification and seed germination analysis of *MeGRX058*-OE, *MeGRX232*-OE, *MeGRX360*-OE and *MeGRX785*-OE *Arabidopsis*.

Supplementary figure S3. Germination analysis of *MeGRX232-OE* transgenic *Arabidopsis* on MS supplement with ABA.

Supplementary figure S4. Phylogenic analysis of TGA factors from *Arabidopsis* and cassava.

Supplementary figure S5. Expression analyses of *MeGRX232* in different tissue from drought stressed cassava cultivar Arg7 and SC124.

Supplementary table S1. Protein sequences of glutaredoxins in *Arabidopsis* and cassava.

Supplementary table S2. RNA-seq data of CC-type GRXs in drought stressed cassava.

Supplementary table S3. List of primers used for qPCR analysis.

Supplementary table S4. GO results of MeGRX232 regulated genes in transgenic *Arabidopsis*.

Supplementary table S5. DNA sequence of *MeGRX232* promoter region.

## Acknowledgements

This work was supported by the National Natural Science Foundation of China (grant no. 31401434) to M.B.R., the National Key Technology R&D Program of China (grant no. 2015BAD15B01), the National Natural Science Foundation of China NSFC-CGIAR Project (grant no. 31561143012) to M. P., and the Hainan province innovative research team foundation (grant no. 2016CXTD).

